# Split selectable marker mediated gene stacking in plants

**DOI:** 10.1101/2022.11.19.517202

**Authors:** Guoliang Yuan, Haiwei Lu, Kuntal De, Md Mahmudul Hassan, Wellington Muchero, Gerald A. Tuskan, Xiaohan Yang

## Abstract

Plant synthetic biology and genetic engineering depend on the controlled expression of transgenes of interest, which relies heavily on the use of *Agrobacterium*-mediated transformation. Here we establish a novel split selectable marker system using protein splicing elements called “inteins” for *Agrobacterium*-mediated co-transformation in plants. This method enables robust co-transformation in *Arabidopsis* and poplar, providing a novel strategy for the simultaneous insertion of multiple genes into both herbaceous and woody plants.

## Main

Metabolic engineering in plants relies on the introduction of complete or partial synthetic pathway into the target plant to create novel plant traits and produce value-added metabolites and therapeutic proteins ^1,2^. A complete synthetic pathway is typically encoded by multiple genes that involves multiple genetic parts and gene circuits. Multigene engineering therefore is becoming more and more important for plant synthetic biology research. Also, a lot of complex plant traits (e.g., yield) are controlled by multiple genes, and as such, genetic improvement of such polygenic traits requires multi-gene stacking. *Agrobacterium*-mediated transformation is to date the most widely used method for plant genetic engineering due to its relatively high efficiency^3^. Although some progress in *Agrobacterium*-mediated transformation of large DNA fragments required for multigene engineering in plants has been achieved, it has been reported that large genomic DNA fragments are not stable in *Agrobacterium* and T-DNA can be truncated at the left and/or right ends before being inserted into the plant genome ^4–6^. Thus, effective transformation of tens of genes into a plant genome and consequent optimal control of gene expression remain unattainable in plant engineering. Here, we developed a split-intein-based gene-stacking method through split-selectable-marker-enabled co-transformation in *Arabidopsis thaliana* and poplar (*Populus*).

An intein is an intervening protein domain that excises itself post-translationally from the host protein leading to the ligation of flanking N- and C-terminal residues (termed as exteins), which share a common intein, to form a new protein (Figure 1a) ^7^. Split inteins to date have enabled the development of numerous tools for both synthetic and biological applications by providing a rapid and bioorthogonal means to link two polypeptides ^7,8^. As such, we previously show that split intein, derived from *Npu*DnaE ^9^, can be used to reduce the size of CRISPR/Cas9 system by split Cas9 into multiple fragments, leading to effective base editing in plants ^10^. In this study, we aim to develop split selectable markers using split inteins to enable single-selectable-marker-gene dependent co-transformation in plants. Initially, we selected eYGFPuv ^11^ and RUBY ^12^ as the reporter genes that can be easily visualized by naked eyes with and without UV light, respectively, to establish a functional split system. In fact, the RUBY reporter is encoded by three genes *CYP76AD1, DODA*, and *glucosyltransferase* (*GT*) (Figure 1b). In general, a catalytic Cys residue at position +1 of the C-terminal extein is mandatory to maintain substantial splicing activities ^7^. A potential split site, L231:C232, was thus identified for splitting the gene *GT* into two fragments, termed GTf1 and GTf2 (Figure 1c). We split the RUBY reporter into two parts by creating plasmid pAXY0006 containing *CYP76AD1, DODA* and GTf1-NpuDnaE(N), and plasmid pAXY0007 containing NpuDnaE(C)-GTf2 (Supplementary Figure 1). Note that the *Arabidopsis* codon-optimized *NpuDnaE* intein was created to improve gene expression and translational efficiency in plants. We then tested split-RUBY using *Agrobacterium*-mediated leaf infiltration in *Nicotiana benthamiana*. Strong red pigment was observed by naked eyes both in the positive control (RUBY) and the leaf area co-infiltrated with pAXY0006 and pAXY0007 plasmids, whereas no red pigment was detected in the leaf area infiltrated with pAXY0006 or pAXY0007 plasmids alone (negative controls) (Figure 1d). Taken together, our results indicate that the functional RUBY reporter was restored by split inteins, which is consistent with the split-eYGFPuv reported previously ^10^.

**Figure 1.**
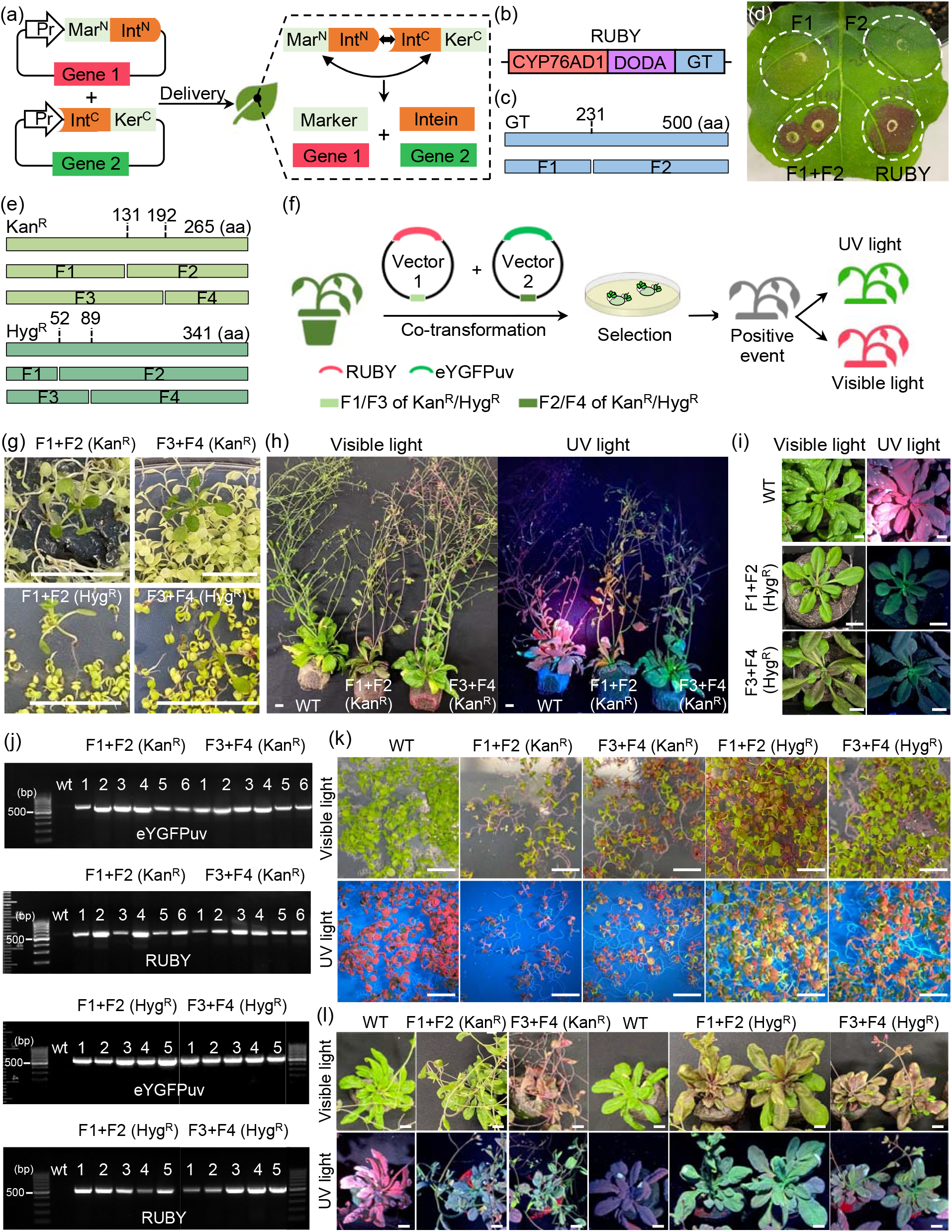
Split RUBY and the split–selectable markers mediated gene stacking in tobacco and *Arabidopsis*. (a) Split selectable marker for co-selection of two separate transgenic vectors. (b) Illustration of RUBY reporter. (c) Identification of potential split site of gene *GT*. (d) Transient expression of split-RUBY in *N. benthamiana*. (e) Identification of potential split sites of gene Kan^R^ and Hyg^R^. (f) Experiment design of the split–selectable markers mediated co-transformation in plants. (g) Selection of T1 seedling on MS medium containing Kanamycin or Hygromycin. (h) Phenotyping of Kanamycin resistant T1 transformants (six weeks). (i) Phenotyping of Hygromycin resistant T1 transformants (four weeks). (j) Genotyping of T1 transformants using primers of eYGFPuv and RUBY, respectively. (k) Selection of T2 seedling on MS medium containing Kanamycin or Hygromycin. (l) Phenotyping of Kanamycin-resistant T2 transformants (five weeks) and Hygromycin-resistant T2 transformants (four weeks). Scale bar, 1 cm.

Because Kanamycin resistance (Kan^R^; *nptII*) and Hygromycin resistance (Hyg^R^; *hpt*) are widely used as the selectable markers in plant transformation, we next tested *NpuDnaE* intein ^9^ for splitting the *nptII* gene encoding neomycin phosphotransferase II and the *hpt* gene encoding Hygromycin phosphotransferase, which confer Kan^R^ and Hyg^R^, respectively. Following the rule of obligatory cysteine residue on the C-extein ^7^, we identified two split sites (T131:C132 and A192:C193) for *nptII* and two split sites (S52:C53 and Y89:C90) for *hpt* (Figure 1e).

The coding sequence of *nptII* or *hpt* was split into an N-terminal fragment (Mar^N^, F1/F3) and a C-terminal fragment (Ker^C^, F2/F4), which were then cloned upstream of an N-terminal fragment of the *NpuDnaE* intein (Int^N^) and downstream of a C-terminal fragment of the *NpuDnaE* intein (Int^C^), respectively, into two vectors (Figure 1a and e). Each vector also carries one of the two reporters (eYGFPuv and RUBY), which allow for easy assessment of co-transformation (Supplementary Figure 2). Thus, we expect to see both green fluorescence under UV light and red pigment under visible light in Kanamycin-resistant or Hygromycin-resistant transgenic plants after co-transformation of these two vectors (Figure 1f).

After co-transformation via floral dip in *Arabidopsis*, multiple transgenic seedlings with typical Kanamycin-resistant or Hygromycin-resistant phenotype were successfully identified on the selection media, indicating that the two inactive fragments of each selectable marker gene (*nptII* or *hpt*) were effectively reconstituted post-translationally (Figure 1g). Subsequently, green fluorescence and red pigment were observed at different stages of Kanamycin-resistant T1 plants (Figure 1h) and Hygromycin-resistant T1 plants (Figure 1i) under UV light and visible light, respectively, suggesting that both eYGFPuv and RUBY vectors were also transformed into the same plant simultaneously through split-Kan^R^- or split-Hyg^R^-mediated co-transformation. These observations were further confirmed by PCR-based genotyping, where both *eYGFPuv* and *RUBY* genes were detected in all Kanamycin-resistant or Hygromycin-resistant plants (Figure 1j). Next, we evaluated the phenotype of T2 generations of above transgenic plants, with the expectation that the Kanamycin-resistance or Hygromycin-resistance phenotype will be observed in T2 seedlings, along with green fluorescence under UV light and red pigment under visible light, as seen in Figure 1k. The phenotypes of eYGFPuv and RUBY were continuously detected in the mature plants of Kanamycin-resistant lines and Hygromycin-resistant lines (Figure 1l and Supplementary Figure 3). These results show that the traits of Kanamycin-resistant, Hygromycin-resistant, eYGFPuv and RUBY are all heritable in *Arabidopsis* across generations, demonstrating the split–Kan^R^-and split–Hyg^R^-mediated co-transformation are effective and robust methods for stable gene-stacking in plants.

We further examined the efficacy of split–Hyg^R^ system in poplar. After tissue-culture-based co-transformation in poplar ‘717’ (*Populus tremula* x *alba* clone INRA ‘717-1B4’), we observed more than 20 transgenic shoots that shows bright green fluorescence under UV light. We randomly selected 15 eYGFPuv-expressing shoots and cultured them on root induction medium supplied with Hygromycin (Figure 2a), where 80% of induced shoots were rooted successfully on selection medium, suggesting that functional Hygromycin phosphotransferase was generated post-translationally. We observed consistent green fluorescence in all rooted plants over time though some plants showed weaker and nonuniform eYGFPuv signals (Figure 2a). Interestingly, typical RUBY phenotype was not observed in the rooted plants though red pigmentation was observed in the stem of some plants. Intriguingly, we detected both eYGFPuv and RUBY genes via PCR genotyping in all rooted plants (Figure 2b and c). Overall, these results suggests that the split–Hyg^R^ system can also work effectively in tissue-culture-based transformation in woody plants. To directly observe protein splicing and to confirm these inteins are indeed orthogonal, we conducted western blot analysis of protein trans-splicing between N-Hyg^R^ N-terminally tagged with 3xFLAG-epitope and C-Hyg^R^ C-terminally tagged with 3xHA-epitope (Figure 2d and Supplementary Figure 4). As expected, full-length Hyg^R^ (lanes 6,7) was observed in the co-transformation of cognate Hyg^R^ fragments with matching N- and C-inteins while transformation with fragment F1/F2/F3/F4 only (lanes 2,3,4,5) did not yield full-length HygroR (Figure 2d).

**Figure 2.**
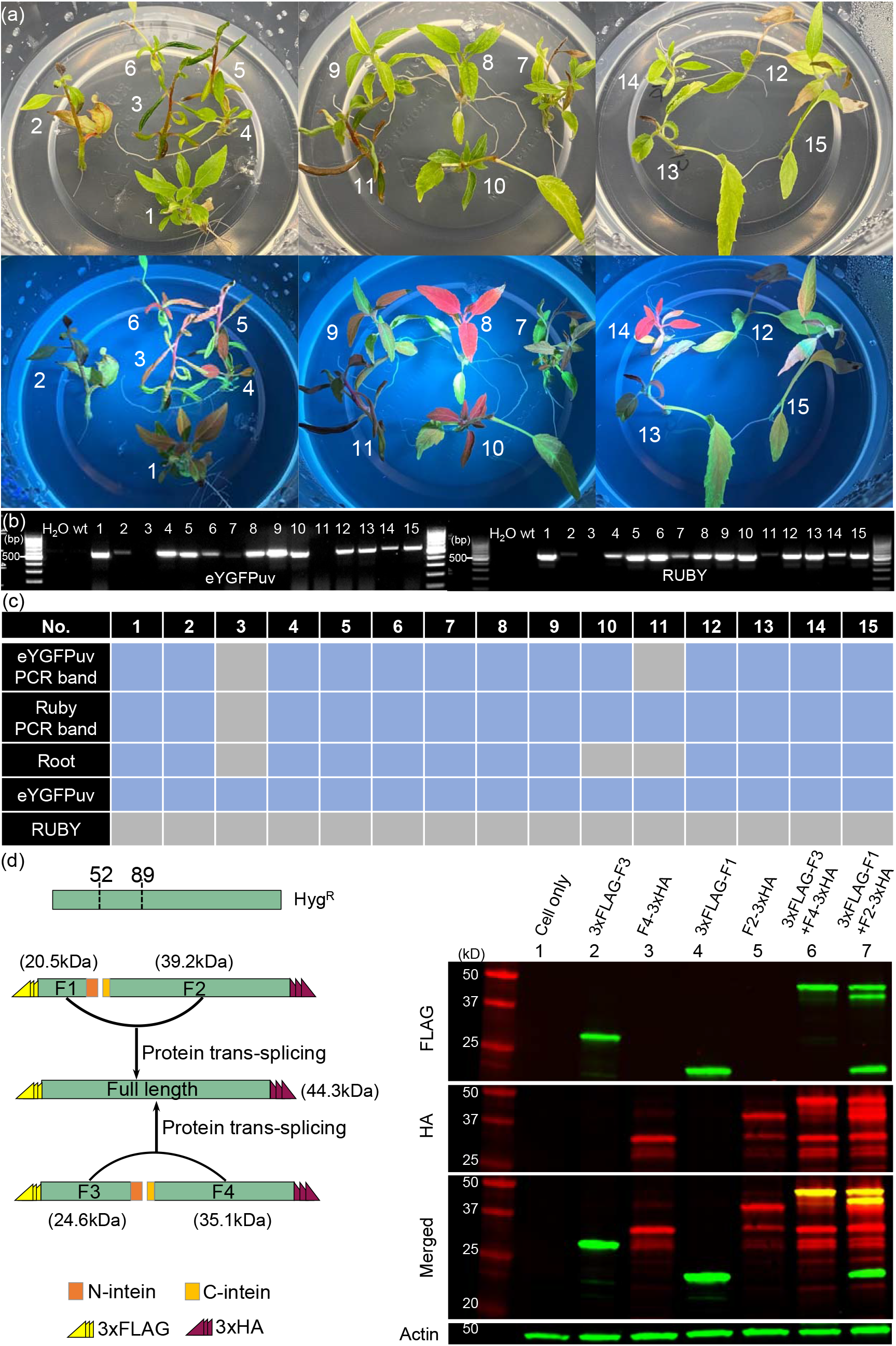
The split–selectable markers mediated gene stacking in poplar. (a) Root induction in root induction medium supplied with Hygromycin and phenotyping of transgenic events with/without UV light. (b) Genotyping of transgenic poplar events using primers of eYGFPuv and RUBY, respectively. (c) The analysis and alignment of genotyping and phenotyping of transgenic plants. Blue block represents positive events while gray block represents negative events. (d) Western blot analysis of trans-splicing of the Hyg^R^ protein. The N-terminal fragments of Hyg^R^ (F1 and F3) are N-terminally tagged with 3xFLAG epitope while the C-terminal fragments of Hyg^R^ (F2 and F4) are C-terminally tagged with 3xHA epitope. Western blot was performed with the proteins extracted from human kidney cells, which were either transfected with the plasmids containing one of fragment F1 to F4, respectively, or co-transfected with F1+F2 containing plasmids and F3+F4 containing plasmids. Red, green, and yellow bands indicate FLAG, HA, and merged bands, respectively. Actin serves as the equal-loading control.

Current plant co-transformation approaches rely on at least two selectable gene markers. For this protocol the concentrations of combined antibiotics need to be tested and adjusted carefully to achieve optimal transgenic selection effect. There is also a difference in selection efficacy between different selectable markers, such that, Hyg^R^ works better (lower rate of false positives) than Kan^R^ in the genetic transformation of some poplar genotypes ^13,14^. In this study, for the first time in plants, we demonstrate that the systems of split–Kan^R^ and split–Hyg^R^ are effective for both *in planta* and plant tissue culture co-transformation in herbaceous and woody plants. This technology potentially doubles the capacity of existing transformation systems for multi-gene engineering in plants. Furthermore, the transgenic events can be easily, visually identified using RUBY and eYGFPuv reporters without a need for expensive advanced resources and labor-intensive process ^11,12,15^. Similarly, the advantages of these co-transformation methods can reduce valuable time spent on constructing complex or long T-DNA molecules in binary vectors and sequential transformations, thus improving the capabilities for pathway engineering and genetic improvement of polygenic traits. Finally, the current common practice of expressing multiple genes involves the repeated use of the same or similar promoters, due to the limited number of available promoters ^16^. Here, repetitive sequences within a plasmid can undergo intramolecular DNA recombination ^17^. This scenario is avoided with the use of split selectable marker system described here. The choice of delivering multiple gene expression cassettes containing multiple identical sequences with two transformation vectors should allow a drastic reduction of the frequency of plasmid DNA recombination.

## Methods

### Plant materials

*Arabidopsis* (*Arabidopsis thaliana*) ecotype Columbia-0 (Col-0) and tobacco (*Nicotiana benthamiana*) were grown in controlled-climate chambers under fluorescent cold white light (100 to 150 μmol m^-2^ s^-1^), 16-h light/8-h dark photoperiod, 20–22 °C, and 60% humidity. *In vitro*–grown poplar ‘717’ (*Populus tremula* x *P. alba* clone INRA 717-1B4) plantlets were placed in a growth room with photoperiod of 16-h light/8-h dark at 22 °C.

### Vector construction

To split RUBY, we first created a RUBY-minus vector lacking the gene *GT* by assembling PCR product 1 containing *CYP76AD1* and *DODA* and PCR product 2 containing *Arabidopsis* HSP18.2 terminator into a pGFPGUSplus vector ^18^ via NEBuilder HiFi DNA Assembly (New England BioLabs). The pAXY0006 vector of split-RUBY was generated by assembling PCR products containing f1 fragment of gene *GT* (named GTf1) and NpuDnaE(N) into RUBY-minus vector via NEBuilder HiFi DNA Assembly. The pAXY0007 vector of split-RUBY was generated by assembling PCR products containing f2 fragment of gene *GT* (named GTf2) and NpuDnaE(C) into pGFPGUSplus vector via NEBuilder HiFi DNA Assembly. To split Kan^R^ (i.e., *nptII*) and Hyg^R^ (i.e., *hpt*), gBlocks Gene Fragments containing either 5’-Kan^R^/Hyg^R^ and N-terminal of NpuDnaE or C-terminal of NpuDnaE and 3’-Kan^R^/Hyg^R^ were synthesized from Integrated DNA Technologies IDT. The pAXY0008/00010/00012/00014 vectors of split-Kan^R^/Hyg^R^ were generated by assembling PCR products containing F1/F3 fragment of Kan^R^/Hyg^R^ and NpuDnaE(N) into pGFPGUSplus vector via NEBuilder HiFi DNA Assembly. The pAXY0009/00011/00013/00015 vectors of split-Kan^R^/Hyg^R^ were generated by assembling PCR products containing F2/F4 fragment of Kan^R^/Hyg^R^ and NpuDnaE(C) into pGFPGUSplus vector via NEBuilder HiFi DNA Assembly. The coding sequences of inteins were codon optimized for *Arabidopsis* via the online codon optimization tool (ExpOptimizer) provided by NovoPro Bioscience (Shanghai, China). All vectors were verified by Sanger sequencing. Information for all primers, gBlocks and plasmids used in this study is provided in Supplementary Table 1 and 2.

### *Arabidopsis* stable transformation

The *Agrobacterium tumefaciens* strain ‘GV3101’ was used for the transformation of *Arabidopsis* wild type ‘Col-0’ via the floral dip method as described by Yuan *et al* ^19^. For co-transformation, two *Agrobacterium* strains containing corresponding vectors (Supplementary Figure 2), respectively, were cultured separately overnight in 100 mL LB liquid medium supplied with 50 mg/L Kanamycin and 50 mg/L Rifampicin. Two LB cultures were spun down at 4000-5000 rpm for 20 mins and resuspended in 30 mL new LB liquid medium without antibiotics and mix equally. Mixed LB culture was added into 120 mL dip solution containing 5% sucrose and 0.03% Silwet-L77. In general, 8 to 12 plants were used for each co-transformation.

### Poplar stable transformation

The *Agrobacterium tumefaciens* strain ‘EHA105’ was used for the co-transformation of the poplar ‘717’ following a published method ^20^. 50 mL LB culture for each *Agrobacterium* strain was prepared and spun down as described above. Two *Agrobacterium* pellets were resuspended equally in MS induction medium containing 20 μM acetosyringone at an OD_600_ nm of 0.5-0.8 for each strain. Excised leaf disks from young leaves (~150) were soaked in *Agrobacterium* solution for 1 h, followed by multiple steps including co-culture, washing, callus induction, shoot induction, shoot elongation, and root induction.

### Tobacco leaf infiltration

Infiltration of tobacco leaf was performed following a published method ^20^. For co-infiltration, 5 mL overnight culture of two *Agrobacterium* strains were spun down and resuspended equally in resuspension solution containing 10 mM MgCl_2_, 10 mM MES-K (pH 5.6), and 100 μM acetosyringone at an OD_600_ nm of 0.5 for each strain.

### Genotyping

To genotype the resistant lines, leaves, approximately 0.5-1.0 cm, were collected from *Arabidopsis* and poplar ‘717’, and ground to a powder. Genomic DNA was isolated by a modified sodium dodecyl sulfate (SDS)-based DNA extraction method. Forward primer 5’-CACGGCAACCTCAACG-3’ and reverse primer 5’-CTCGACACGTCTGTGGG-3’ were used for genotyping PCR of *eYGFPuv*. Forward primer 5’-CAGAGCTTGCGAGAAAGG-3’ and reverse primer 5’ - GGCGGAGGTGAACTTGTAG-3’ were used for genotyping PCR of *RUBY*.

### Phenotyping

The fluorescence signals of eYGFPuv were visualized under a 365 nm wave-length UV light and imaged using an iPhone 11 as described by Yuan *et al*^11^. The red pigment due to RUBY expression is visible by naked eyes without requiring any equipment ^12^ and images were also taken using an iPhone 11.

### Protein extraction and Western blot

HEK 293T cells were obtained from ATCC and maintained in a humidified atmosphere at 5% CO_2_ in Dulbecco’s Modified Eagle’s (DMEM) complete medium (Corning) supplemented with 10% fetal bovine serum (FBS; Seradigm) in 37 °C. Plasmid transfections were done with TransIT-LT1 (Mirus Bio) per the manufacturer’s instructions. Briefly, cell extracts were generated on ice in EBC buffer, 50 mM Tris (pH 8.0), 120 mM NaCl, 0.5% NP40, 1 mM DTT, and protease and phosphatase inhibitors tablets (Thermo Fisher Scientific). Extracted proteins were quantified using the PierceTM BCA Protein assay kit (Thermo Fisher). Proteins were separated by SDS acrylamide gel electrophoresis and transferred to IMMOBILON-FL 26 PVDF membrane (Millipore) probed with the indicated antibodies and visualized either by chemiluminescence (according to the manufacturer’s instructions) or using a LiCor Odyssey infrared imaging system. Western blot were conducted as described previously ^10^.

## Author contributions

G.Y. and X.Y conceived the research. G.Y., H.L., K.D. and M.H. conducted the experiments. G.Y. wrote the paper. All authors revised the manuscript. X.Y. and G.A.T. supervised the research.

## Acknowledgements

The work was supported by the Center for Bioenergy Innovation, a U.S. Department of Energy (DOE) Bioenergy Research Center supported by the Biological and Environmental Research (BER) program. Oak Ridge National Laboratory is managed by UT-Battelle, LLC for the U.S. Department of Energy under Contract Number DE-AC05-00OR22725.

## Data availability

The plasmids will be available at Addgene.

## Supplementary Information

**Supplementary Figure 1.**
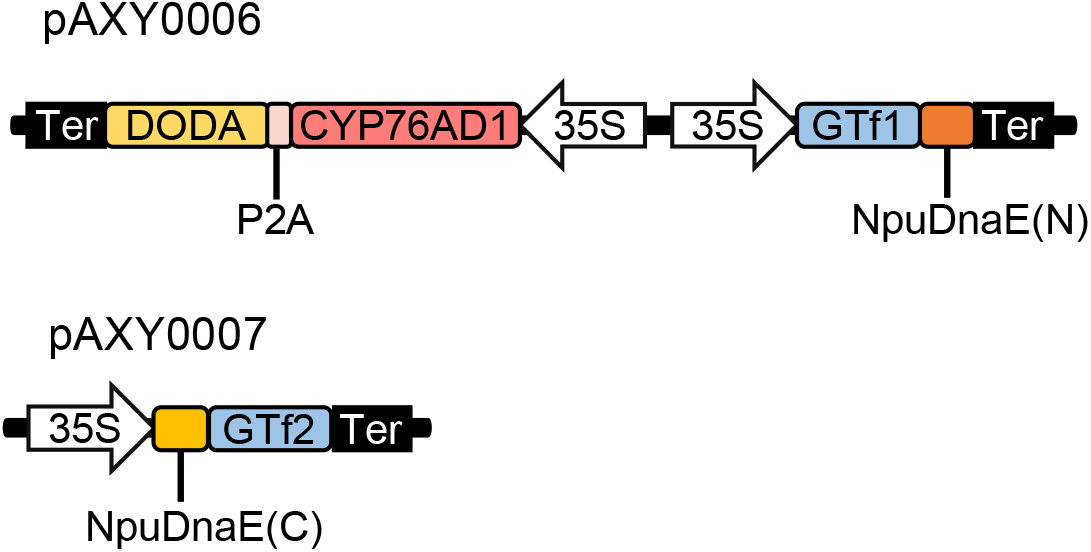
Illustration of split-Kan^R^ and -Hyg^R^ vectors.

**Supplementary Figure 2.**
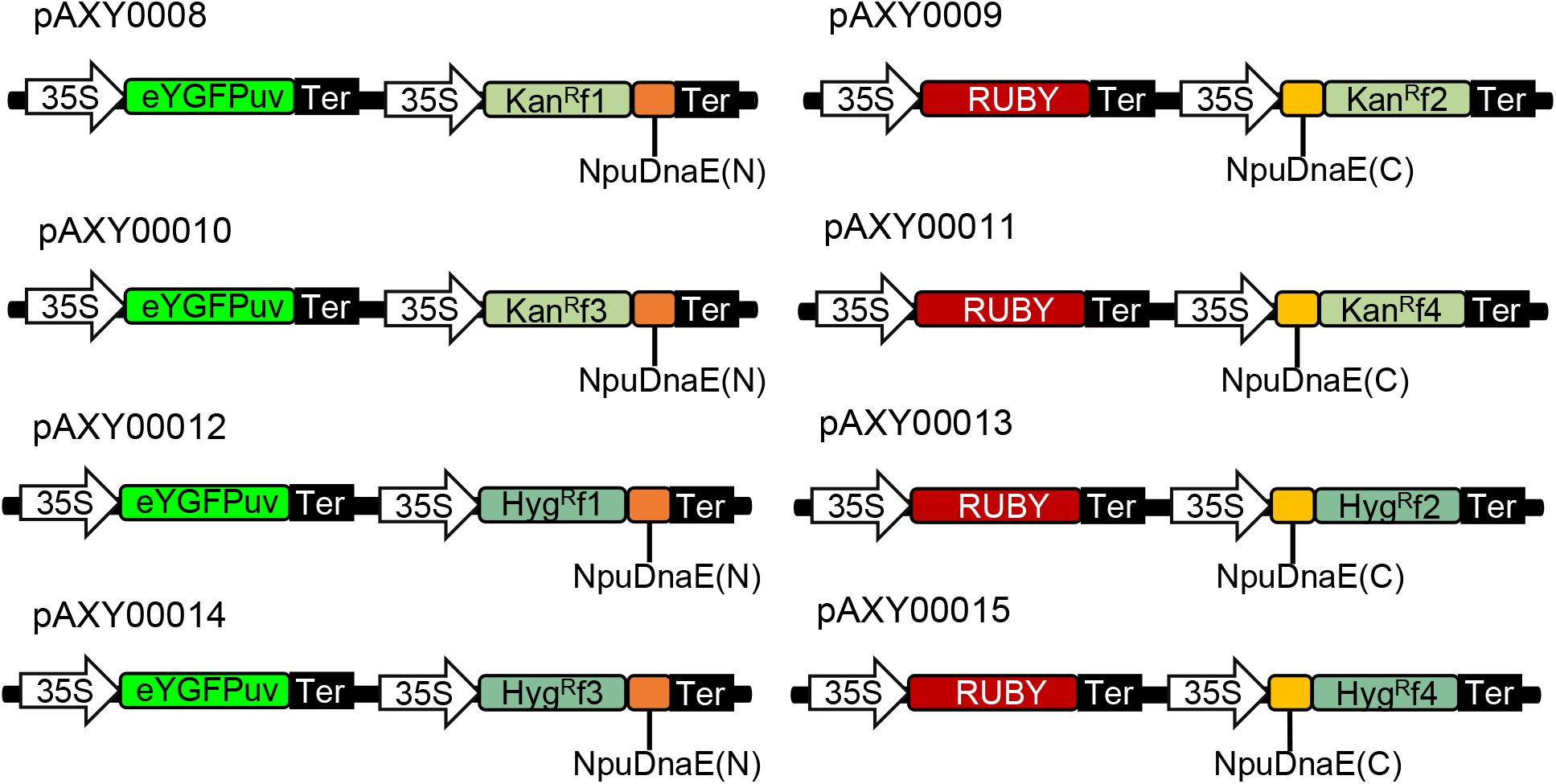
Phenotyping of Kanamycin-resistant and Hygromycin-resisted T2 transformants.

**Supplementary Figure 3.**
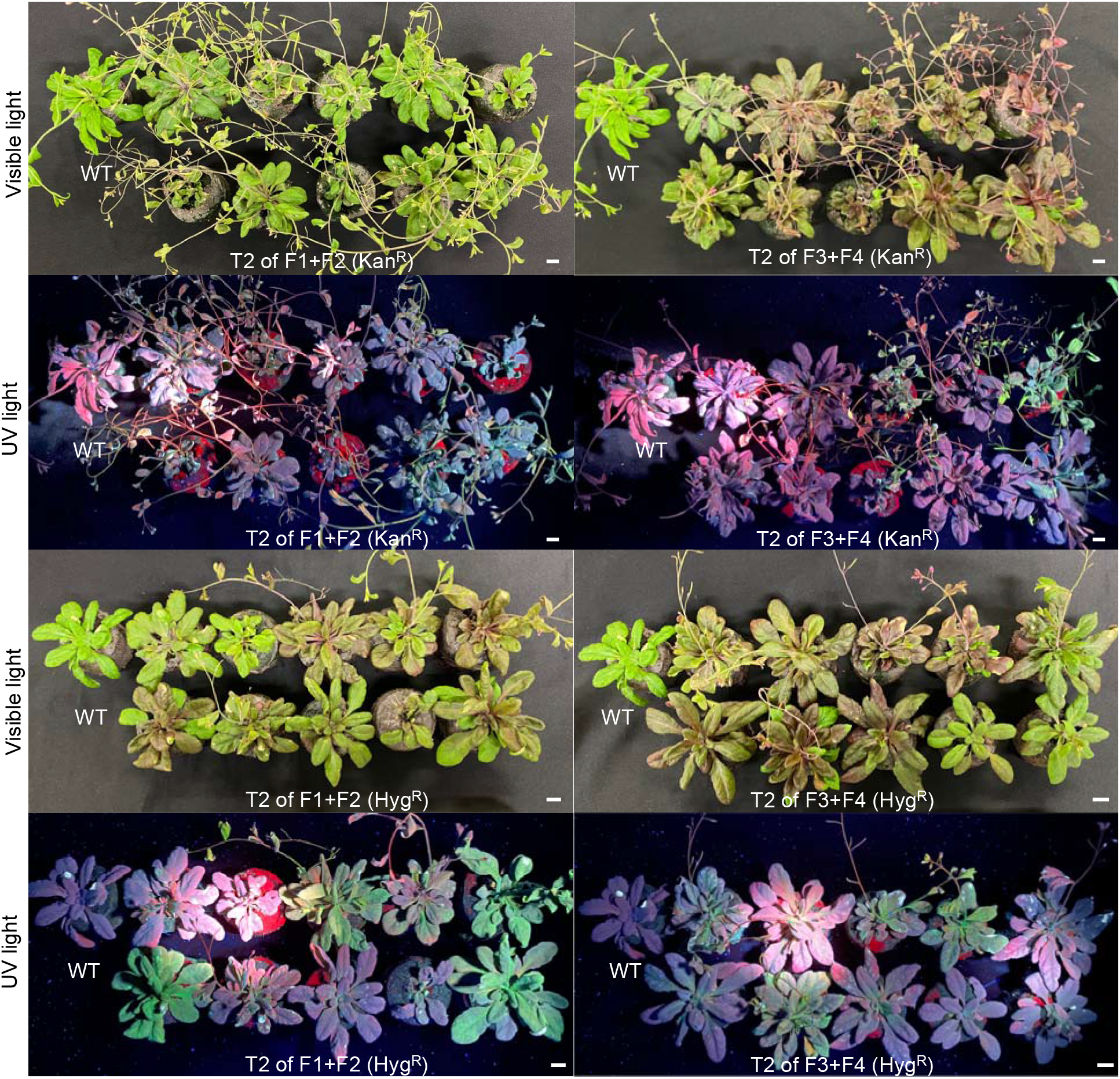
Phenotyping of Kanamycin-resistant and Hygromycin-resisted T2 transformants.

**Supplementary Figure 4.**
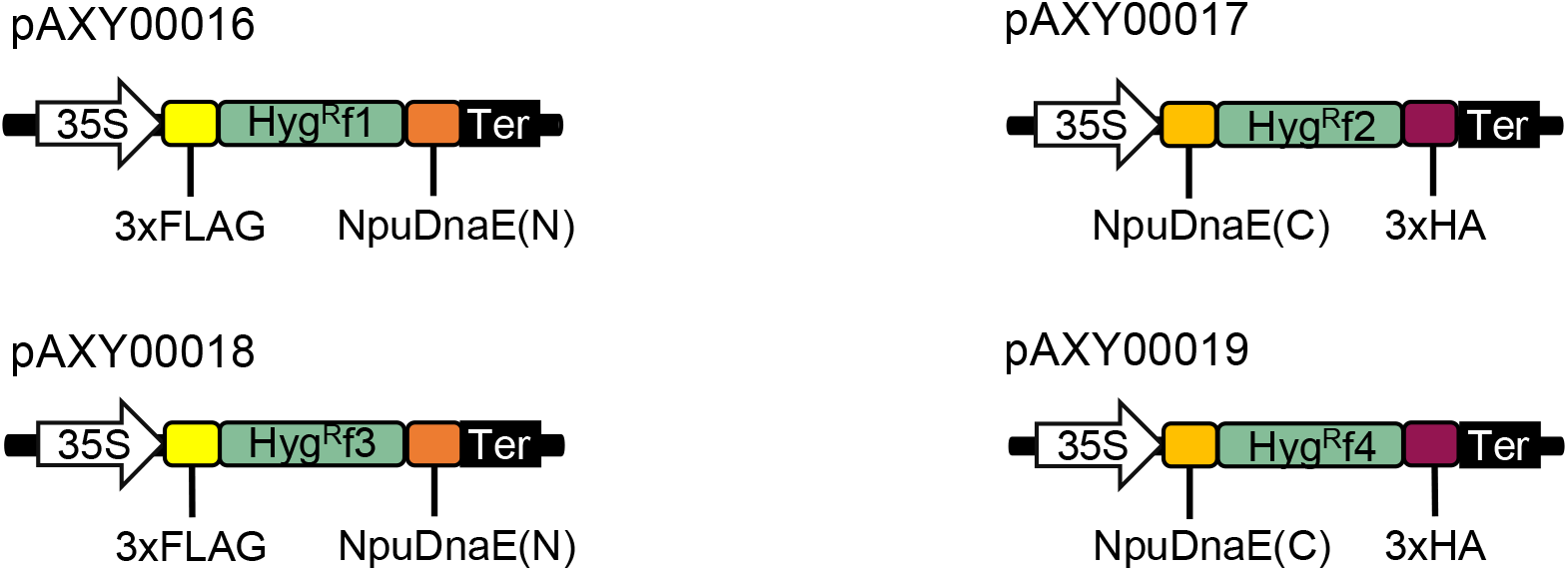
Illustration of split-Hyg^R^ vectors with FLAG or HA epitope.

**Supplementary Table 1.**
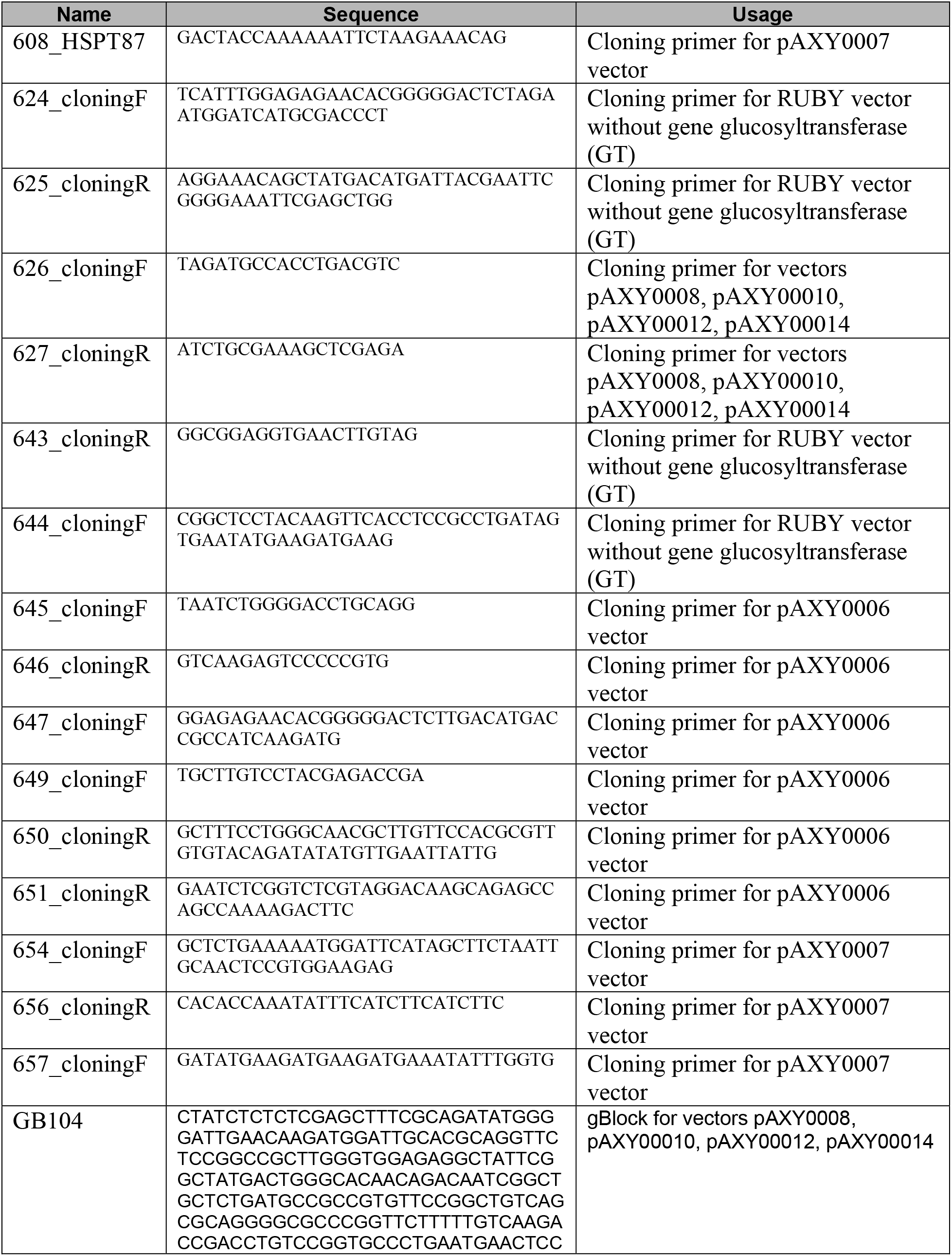
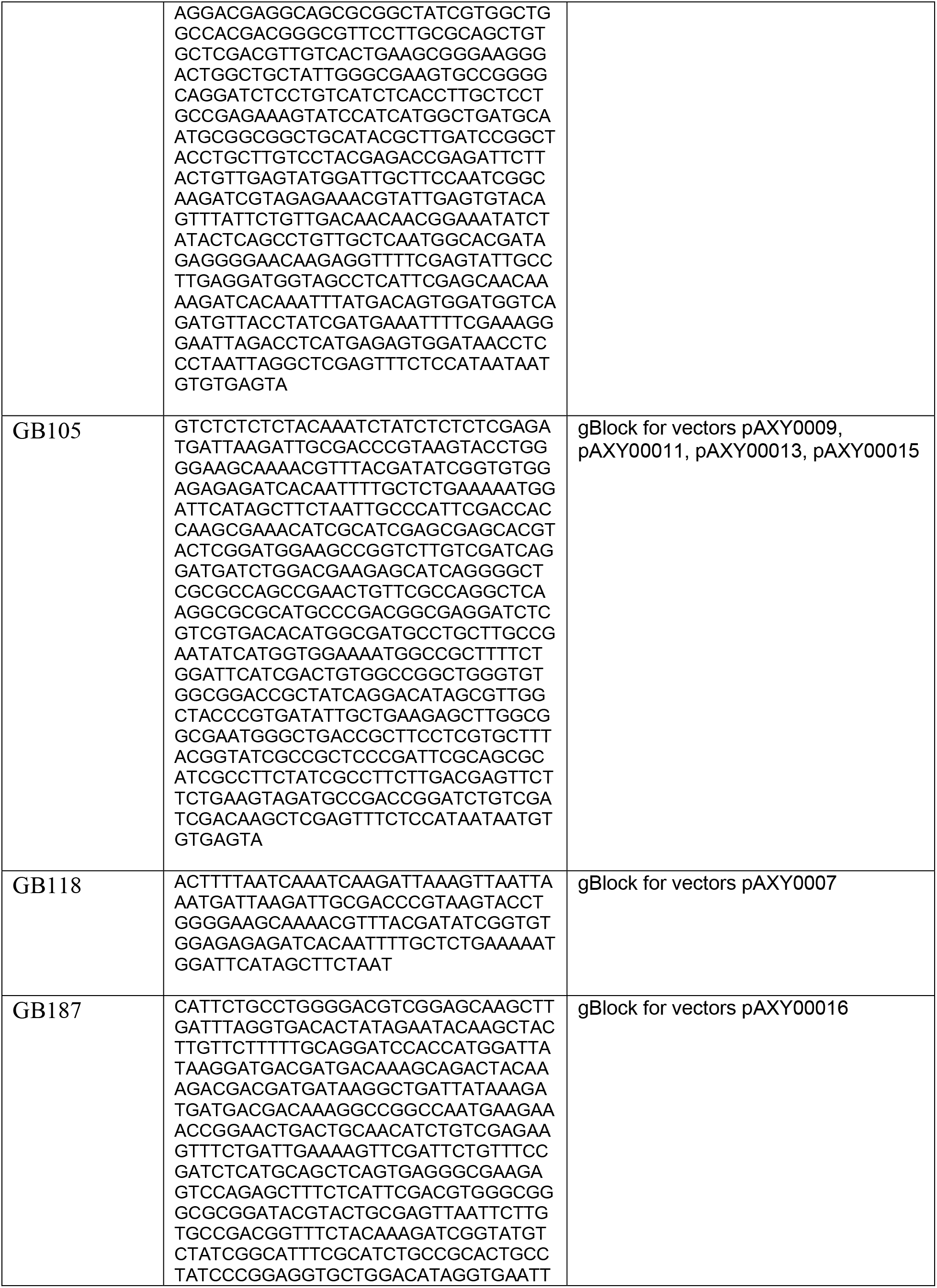
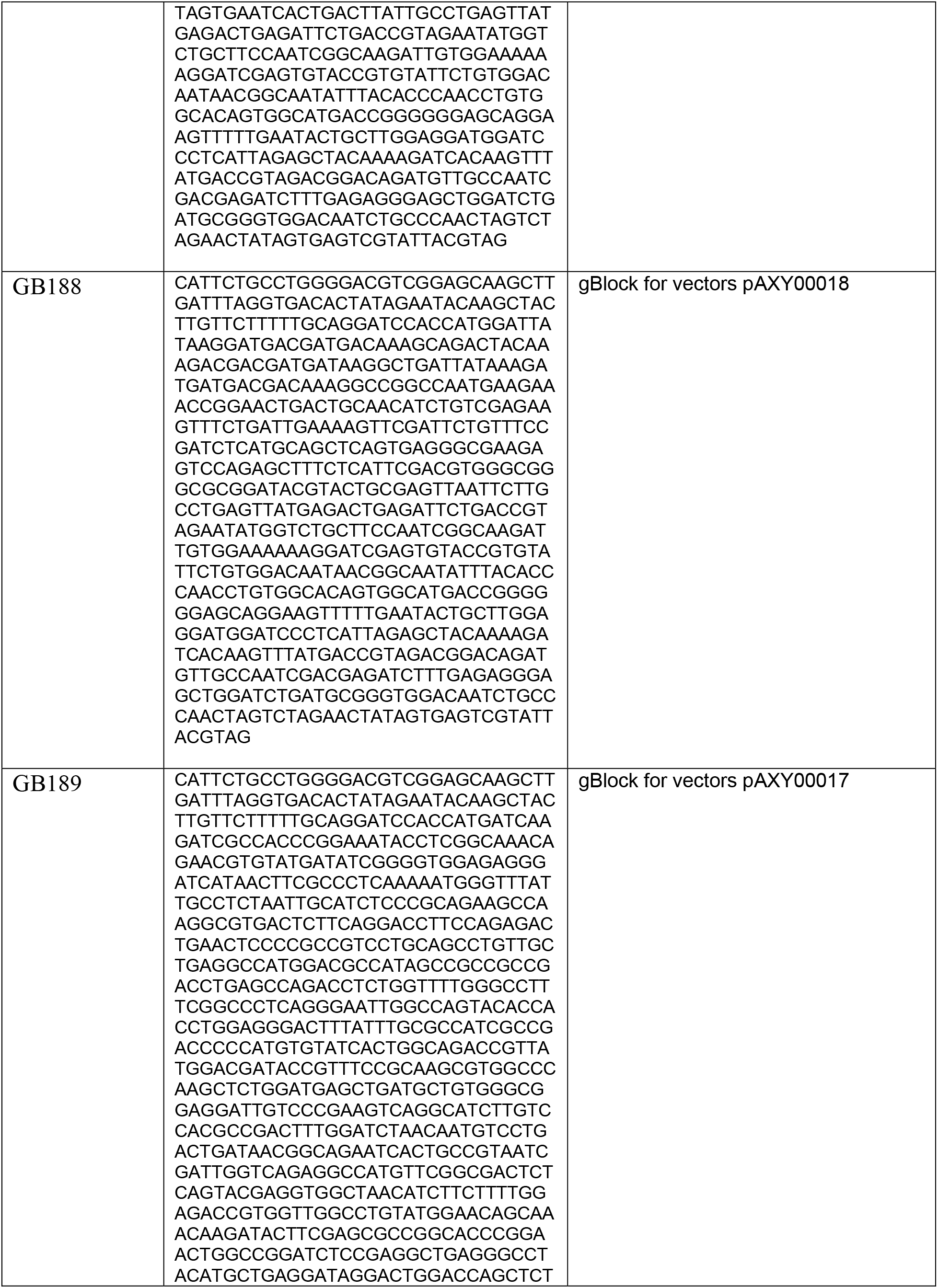
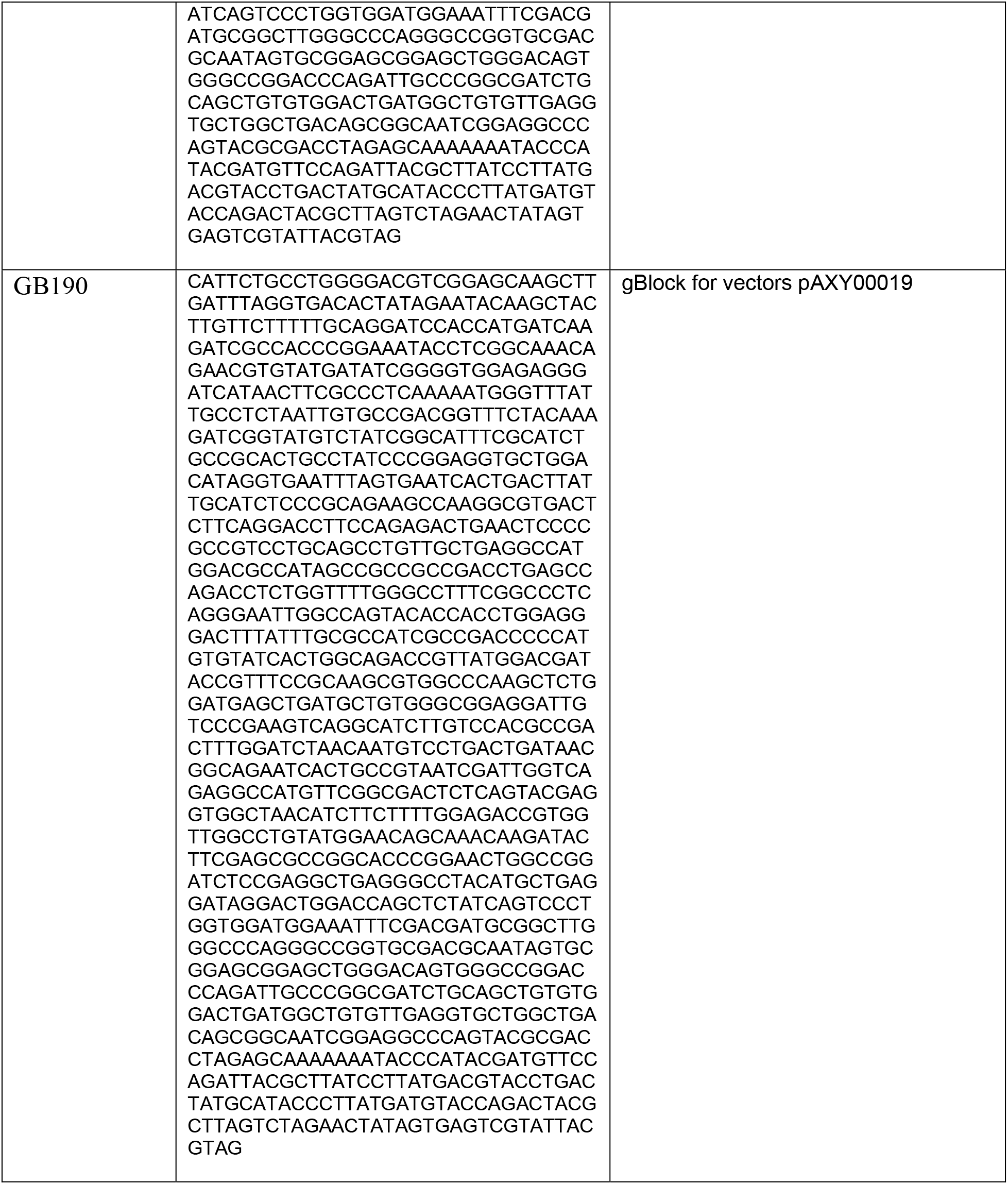
Oligos and gBlocks used in this study.

**Supplementary Table 2.** Plasmids used in this study.

| Plasmid # | pXYA# | Plasmid Name |
| --- | --- | --- |
| 1 | pAXY0006 | pGlucos(1-231)-NpuDnaE(N) |
| 2 | pAXY0007 | pNpuDnaE(C)-Glucos(232-500) |
| 3 | pAXY0008 | pKan<sup>R</sup>(1-131)-NpuDnaE(N) eYGFPuv. |
| 4 | pAXY0009 | pNpuDnaE(C) Kan<sup>R</sup>(132-265) RUBY. |
| 5 | pAXY0010 | pKan<sup>R</sup>(1-192)-NpuDnaE(N) eYGFPuv. |
| 6 | pAXY0011 | pNpuDnaE(C) Kan<sup>R</sup>(193-265) RUBY. |
| 7 | pAXY0012 | pHyg<sup>R</sup>(1-52)-NpuDnaE(N) eYGFPuv |
| 8 | pAXY0013 | pNpuDnaE(C) Hyg<sup>R</sup>(53-341) RUBY |
| 9 | pAXY0014 | pHyg<sup>R</sup>(1-89)-NpuDnaE(N) eYGFPuv |
| 10 | pAXY0015 | pNpuDnaE(C) Hyg<sup>R</sup>(90-341) RUBY |
| 11 | pAXY0016 | p3flag Hyg<sup>R</sup>(1-89)-NpuDnaE(N) |
| 12 | pAXY0017 | pNpuDnaE(C) Hyg<sup>R</sup>(90-341) 3HA |
| 13 | pAXY0018 | p3flag Hyg<sup>R</sup>(1-52)-NpuDnaE(N) |
| 14 | pAXY0019 | pNpuDnaE(C) Hyg<sup>R</sup>(53-341) 3HA |

